# It’s all relative: population estimates enhance kin recognition in the guppy

**DOI:** 10.1101/2020.06.22.165340

**Authors:** Mitchel J. Daniel

## Abstract

Kin recognition plays a fundamental role in social evolution, enabling active inbreeding avoidance, nepotism, and promoting cooperative social organization. Many organisms recognize kin based on phenotypic similarity – a process called phenotype matching – by comparing information associated with their own phenotype against the phenotypes of conspecifics. However, recent theory demonstrates that to accurately judge phenotypic similarity (and hence, relatedness), individuals require estimates of the population’s distribution of phenotypes as a “frame of reference.” Here, I use the Trinidadian guppy (*Poecilia reticulata*) to provide the first empirical test of this population estimation theory. I varied the phenotypic distributions of the groups in which focal individuals developed and found that, as adults, their patterns of inbreeding avoidance and nepotistic intrasexual competition differed as predicted by population estimation theory. Individuals reared with conspecifics more similar to themselves treated novel conspecifics as less closely related, suggesting a shifted population estimate. Individuals reared with more phenotypically variable conspecifics exhibited less extreme kin discrimination, suggesting a broader population estimate. These results provide experimental evidence that population estimates inform phenotype matching, and are acquired plastically through social experience. By calibrating phenotype matching to the population distribution of phenotypes, population estimation enhances kin recognition, increasing opportunities for the evolution of inbreeding avoidance and nepotism.

## Introduction

Social interactions are pervasive in nature, and can have important consequences for fitness. Kin recognition systems allow organisms to assess their genetic relatedness to conspecifics, enabling or enhancing behavioral strategies that increase the fitness consequences of social interactions. Such strategies include active inbreeding or outbreeding avoidance (1), the selective allocation of help towards kin (i.e. nepotism) (2– 4). Kin recognition can also play a fundamental role in biological organization, regulating the formation and maintenance of cooperative kin groups including coalitions (5), social and eusocial societies (6–9), and multicellular bodies as in social amoeba (10–12). While kin recognition is both common and taxonomically widespread (13, 14), the underlying mechanisms remain poorly understood. Resolving these mechanisms remains a key obstacle to answering fundamental questions about the evolution and ontogeny of kin recognition, and the patterns of kin discrimination that occur in nature.

One common and versatile type of kin recognition system is phenotype matching, in which an individual recognizes its kin based on their phenotypic similarity to itself (13, 14). To do so, the evaluator acquires information associated with its own phenotype, either through self-inspection or by observing the phenotypes of its putative kin (e.g. parents or broodmates), and stores this information internally as a “kin template.” Subsequently, the evaluator can compare the kin template against the phenotype of a given conspecific to infer its phenotypic (and hence, genetic) similarity. The evaluator infers relatedness based on the degree of similarity between the conspecific and kin template, and conditions its social actions upon this information. Phenotype matching may, in principle, rely on any phenotypic cue(s)–whether heritable and/or environmentally determined–that closely co-vary with differences in relatedness. The cues involved vary widely among species, and can include olfactory (15–17), visual (18, 19), and acoustic traits (20).

This conceptual model of phenotype matching has largely been taken for granted for nearly 40 years of empirical and theoretical studies of kin recognition (13, 14). However, recent theory suggests that this model is not sufficient to explain how individuals recognize kin (21). Consider a hypothetical case of an evaluator attempting to assess her relatedness to a conspecific. For simplicity, assume she relies on a single continuous phenotype (e.g. height) and uses self-inspection to form the kin template. The evaluator infers her own phenotype (from her kin template) as 0.65 m, and the conspecific’s phenotype as 0.45 m. Under the classical model of phenotype matching, the evaluator’s relatedness judgment should reflect the similarity between these two phenotypes; but, how similar are they? The question cannot be answered with the available information, because an appropriate “frame of reference” is lacking. In addition to knowing its own phenotypes and the conspecific’s phenotype, the evaluator must contextualize the two values using information about the population’s distribution of phenotypes (21).

Information about both the mean and the variance of the distribution of phenotypes are theorized to shape relatedness judgments in important ways (Figure 1). If the evaluator’s phenotype is closer to the conspecific’s phenotype than to the population mean, the two phenotypes are similar and the conspecific is probably a close relative. Alternatively, if the evaluator’s phenotype is further from the conspecific’s phenotype than from the population mean, the two phenotypes are dissimilar, and the conspecific is likely a distant relative. Likewise, the degree of similarity or dissimilarity between the two phenotypes scales inversely with the degree of phenotypic variability in the population. For example, if the phenotype is highly variable, then a small difference between the phenotype of the evaluator and the conspecific implies that they are closely related; conversely, if the population has low phenotypic variation, then even a small difference in phenotype implies that they are not closely related. Thus, phenotypic similarity is intrinsically dependent on the “phenotypic space” of the population, including both the mean and variance in phenotype. By consequence, phenotype matching should only be a reliable means of inferring relatedness if it is informed by some estimate of the population’s distribution of phenotypes.

**Figure 1.**
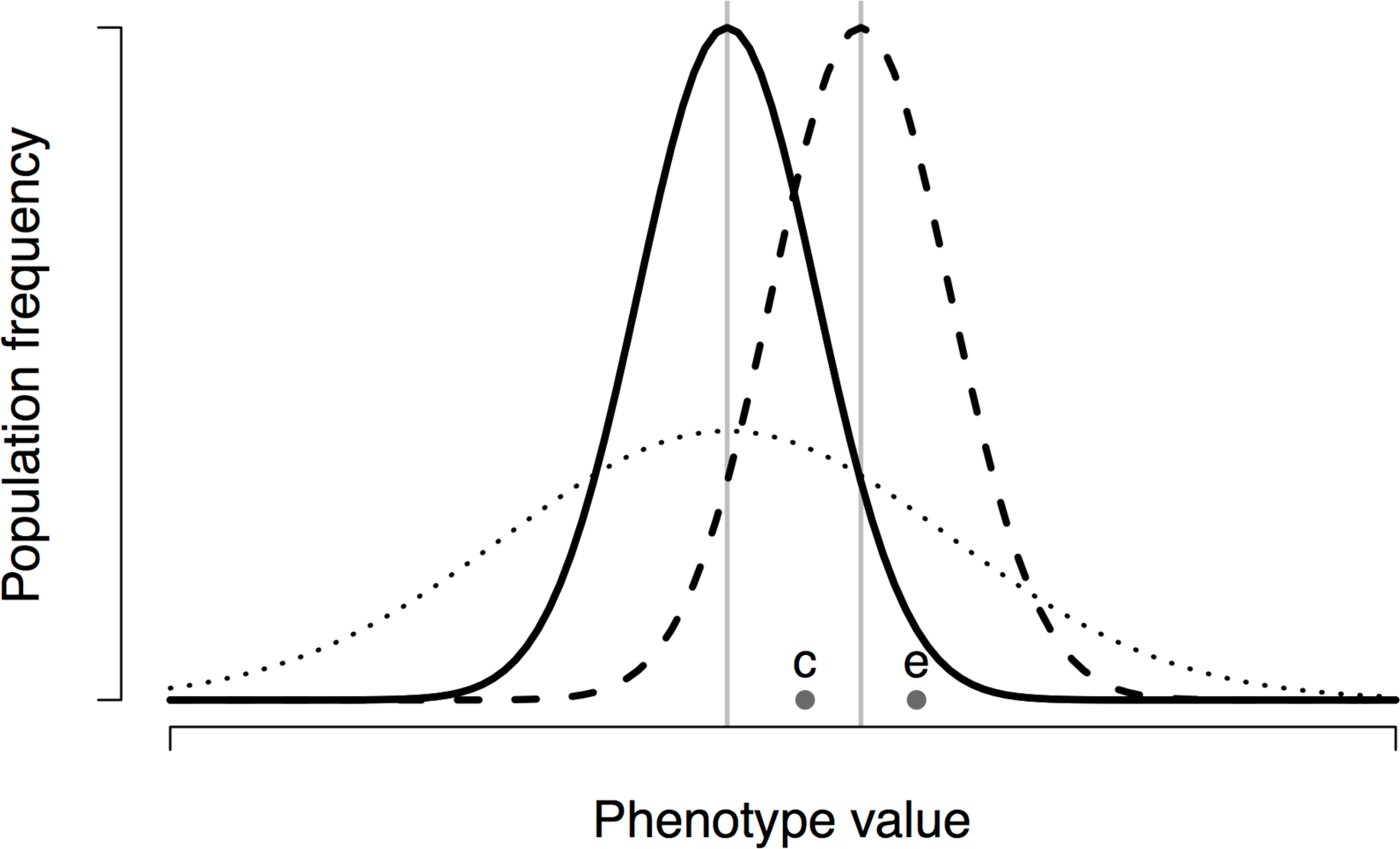
Hypothetical phenotypic distributions with different means (solid vs. dashed) or different variances (solid vs. dotted). Population mean phenotypes are shown with light grey vertical lines. For an evaluator with phenotype e, the phenotypic similarity of a conspecific with phenotype c, differs depending on the phenotypic distribution of their populations. If these two individuals inhabit a population with the phenotypic distribution shown by the solid line, the conspecific’s phenotype is closer to the evaluator’s phenotype than the population average is. Therefore, the conspecific is phenotypically similar and the evaluator should perceive the conspecific as a positive relative (i.e. more genetically similar than average). In contrast, if the same individuals inhabit a population with the phenotypic distribution shown by the dashed line, the conspecific’s phenotype is further from the evaluator’s phenotype than average. The conspecific is, therefore, phenotypically dissimilar and the evaluator should perceive the conspecific as a negative relative (i.e. less genetically similar than average). Furthermore, if the individuals inhabit a population with greater phenotypic variance (solid vs. dotted distributions), the phenotypic space of the population has expanded, such that relatedness judgments should be less extreme (i.e. lower in magnitude, but not different in sign). Thus, even while holding the phenotypes of an evaluator and a conspecific fixed, changes in the population distribution can dramatically alter the appropriate relatedness judgment.

The importance of population estimation to the accuracy of phenotype matching may seem elementary; however, it has only recently gained theoretical attention (21, 22). Moreover, I have not been able to locate any empirical studies investigating whether or how population estimates function in kin recognition. Here, I provide an experimental test of whether individuals use population estimates to enhance kin recognition, and how these estimates are formed.

Population estimates could arise in three ways. First, they could be inherited. At its simplest, inheritance of population estimates could involve a gene that modulates how the evaluator adjusts its social behavior in responses to the kin template and assessment of the conspecific’s phenotype. In this scenario, the population estimate is implicit in the behavioral effects of the gene, and selection should favour whichever allele encodes the most accurate implicit estimate. Second, population estimates could be acquired plastically. For example, individuals may sample and store information about the phenotypes of encountered conspecifics. Third, a population estimate could be both inherited and modified through experience, possibly resembling a Bayesian-like process in which an innate “prior” is updated through learning. Plastic population estimates should confer the advantage of allowing individuals to track changes in the phenotypic distribution over shorter timescales than if population estimates are inherited. Thus, I expect plasticity to be important in systems in which the phenotypic distribution changes relatively quickly through time, as could result from fluctuating population demography or selection on the phenotypes used in kin recognition (23–25). Plasticity should also be advantageous if the distribution of phenotypes varies through space and some individuals disperse to new local populations.

I used the Trinidadian guppy (*Poecilia reticulata*) as a model system for investigating population estimation, a species in which dispersal (26) and fluctuating demography (27–29) should favour the use of plastic population estimates. Phenotype matching is used by female guppies to avoid inbreeding (30), and by juveniles to shoal preferentially with kin (31). Additionally, male guppies use phenotype matching to engage in nepotism during intrasexual competition, directing less competitive effort towards closely related rival males than non-kin rivals (4). Guppies therefore provide an opportunity to investigate the potential role of population estimation in a species in which kin recognition has important fitness consequences in multiple behavioral contexts.

I tested for plastic population estimation by rearing focal individuals within groups that differed either in terms of how similar the mean phenotype was to the phenotype of the focal individual, or in terms of phenotypic variance. I then assayed the patterns of kin discrimination exhibited by these focal individuals, as adults, towards conspecifics spanning a standardized range of phenotypes. I examined patterns of kin discrimination in the contexts of female inbreeding avoidance and male-male competition. I compared the patterns of kin discrimination among individuals reared within different phenotypic distributions to ask (i) whether population estimates play their proposed role in informing phenotype matching, and (ii) whether these estimates are acquired plastically through social experience.

## Methods

### Study system

The guppies in this experiment were lab-reared descendants of the ‘Houde’ tributary of the Paria River in Trinidad (GPS: PS 896 886). The ecology of this population should make plastic population estimates particularly advantageous. This “low-predation” population inhabits an upstream location with pool-and-riffle ecology (27). The spatial kin structure of these upstream locations suggests that pools can be thought of as semi-isolated, local populations linked by dispersal (26). Since phenotype matching relies on phenotypes that co-vary with relatedness, individuals that disperse to a new local population may face a phenotypic distribution different from that of their previous pool. Additionally, seasonal variation in rates of reproduction (27–29, 32) results in fluctuating demography (and hence, changing phenotypic distributions) within pools over time. Given this likely spatial and temporal heterogeneity in phenotypic distributions, I predicted that plastic population estimates enhance kin recognition.

### Background on relatedness and genetic similarity

Kin recognition systems allow individuals to assess their relatedness to conspecifics and adjust their social behavior accordingly. Relatedness itself, *r*, is a measure of differences in genetic similarity, reflecting the degree to which two individuals are more, or less, genetically similar than average within their local population (22, 33, 34). As such, an evaluator’s relatedness to a conspecific depends on their coefficient of consanguinity, *G* (i.e. the probability that both individuals share copies of a randomly selected allele by recent, common descent), and on the evaluator’s mean consanguinity, 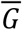, to all conspecifics in the local population (21, 35–37). This formulation accounts for both primary (i.e. actor-recipient) and secondary (i.e. local competition) effects of a social action within the local population (38).

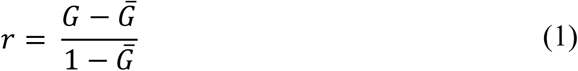

So defined, *r* can take on both positive and negative values reflecting whether a conspecific is more (*r* > 0) or less (*r* < 0) likely to share copies of a given allele with the evaluator than an average member of the population. This metric intuitively describes when, and how, evaluators should adjust their social actions. An individual can increase its indirect fitness by helping positive relatives and harming negative relatives, promoting the evolution of nepotism and spite, respectively (34, 39). Furthermore, inbreeding (or outbreeding) avoidance can be achieved by discriminating against potential mates that are too positively (or too negatively) related (1).

### Experimental design

Previous work suggests that guppies from my study population use olfactory cues for phenotype matching, and acquire the kin template based on self-referencing (i.e. assessment of their own phenotype) (17). Because the specific odor components involved remain unclear, it was not possible, in the present study, to manipulate the distribution of phenotypes directly. Instead, I did so indirectly by varying the *G* (genetic similarity) of focal individuals to the conspecifics within their social group. This is an effective way to indirectly manipulate phenotypic distributions because guppies use heritable phenotypic variation to recognize kin (4, 17, 30). Furthermore, it is likely that variation in the distribution of phenotypes in natural settings is often driven by changes in the distribution of *G* (e.g. demographic heterogeneity), making this approach an ecologically relevant manipulation. Thus, *G* can be thought of as a proxy for phenotypic similarity in my experimental design.

I designed the experiment to test two key predictions made by population estimation theory (21). First, the closer the population mean phenotype is to the evaluator’s phenotype, the less closely related the evaluator should perceive conspecifics with a given phenotype to be. This is because, even though the conspecific’s phenotype is fixed, it should seem less similar to the evaluator’s phenotype by comparison to the population mean (Figure 1). To test this prediction, I reared each focal individual within a “population” consisting of 16 conspecifics. I included a control treatment that served as a baseline for my comparisons, in which the focal individual’s mean *G* to its group members was 0.125, and a high similarity treatment, in which the focal individual’s mean *G* to its group members was higher (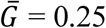; Figure 2A). The distribution of pairwise consanguinities among group members (excluding the focal individual) was the same in both the control and high similarity treatments (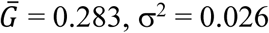) (Figure 2C). Therefore, variance in *G* was equivalent in both the control and high similarity treatments. In this way, I was able to disentangle any effects of mean phenotype and phenotypic variance on population estimation. I predicted that when focal individuals were tested with conspecifics from across the same range of consanguinities to the focal individual, individuals from the high similarity treatment should perceive the conspecifics to be less closely related than should focal individuals from the control treatment (Figure 2E). Focal males from the high similarity treatment should therefore direct more competitive behaviors towards these conspecifics than focal males from the control treatment. Focal females from the high similarity treatment should exhibit greater mating interest in these conspecifics than focal females from the control treatment.

**Figure 2.**
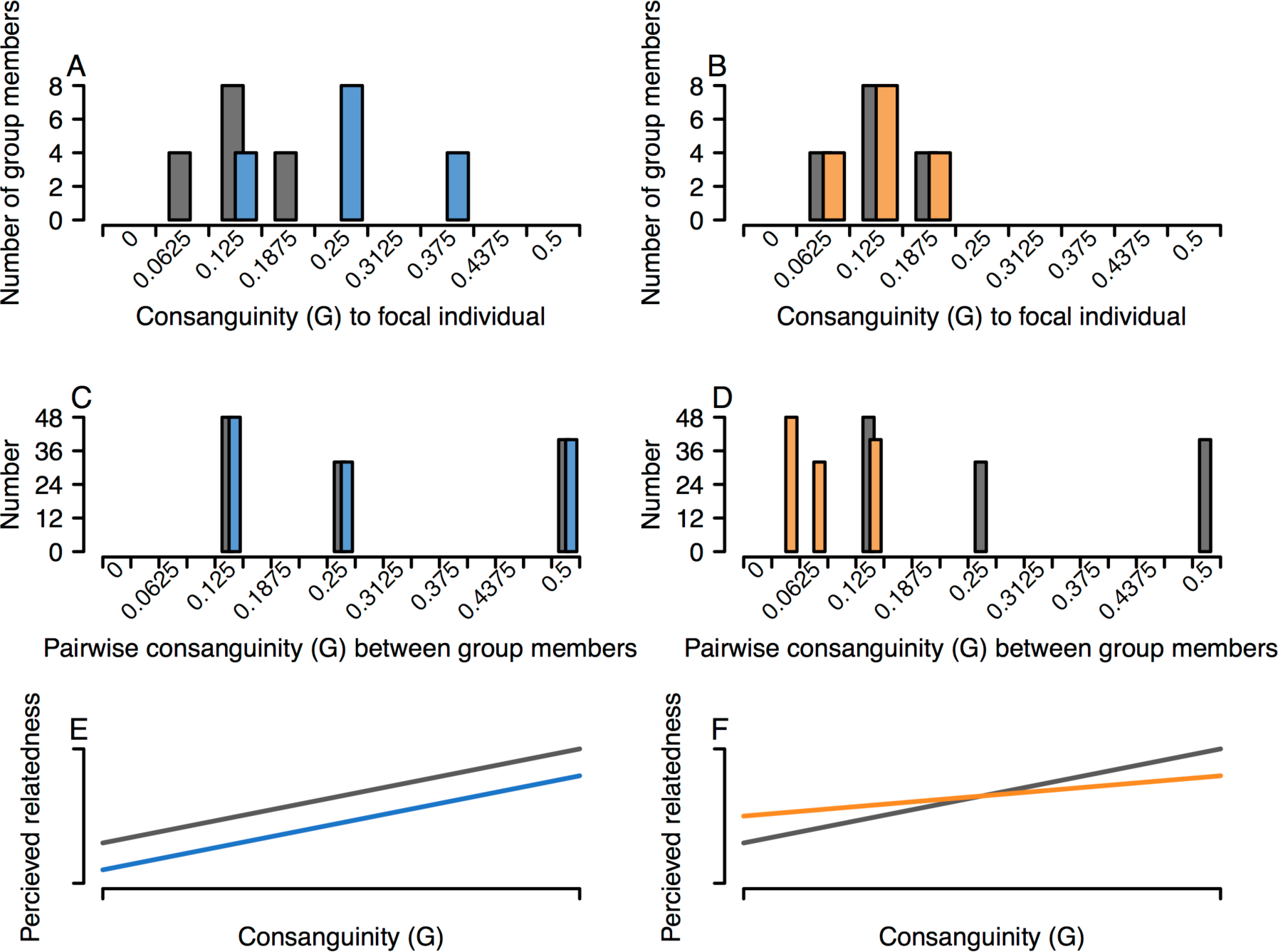
*A,B*: Distribution of consanguinities (*G*) between the focal individual and members of its rearing group in the control (white), high similarity (blue), and high variance (red) treatments. The histograms are offset along the x-axis to improve visibility. *C,D*: Distribution of pairwise *G* between group members for each treatment. *E,F*: Predictions for each treatment, based on population estimation theory, of the focal individuals’ perceived relatedness to a conspecific as a function of *G* to the conspecific. Compared to the control (black) treatment, focal individuals from the high similarity (blue) treatment should perceive conspecifics to be less closely related to themselves. Focal individuals from the high variance (red) treatment should make less extreme relatedness judgments compared to the control. The behaviors I assayed (female mating preference behaviors, and male interruption behaviors) are performed less frequently towards conspecifics that are perceived to be close relatives. Therefore, perceived relatedness should be inversely related to the amount of the behaviors that I measured.

The second prediction I tested is that greater population variance in phenotype should result in the evaluator acquiring a wider frame of reference (Figure 1), resulting in less extreme relatedness judgments. I tested this prediction by comparing the behaviors of focal individuals reared in the control treatment, in which the mean pairwise *G* among group members was 0.283, with that of focal individuals reared in a high variance treatment, in which pairwise *G* among group members was lower (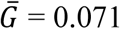; Figure 2D). The distribution of *G* values between the focal individual and its group members was identical for both the control and high variance treatments (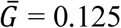, σ^2^ = 0.002; Figure 2B), allowing me to disentangle the effects of phenotype variance from mean phenotype. I predicted that focal individuals from the higher variance treatment should exhibit less extreme relatedness judgments than focal individuals from the control treatment. That is, when focal individuals interact with conspecifics across a given range *G*, the slope of the relationship between *G* and perceived relatedness should be shallower for individuals from more variable groups (Figure 2F).

While the levels of *G* in these distributions may seem high, many low-predation guppy populations exhibit small effective population sizes and local inbreeding (26, 40, 41). Consequently, the population distributions I used should be within the range of some local populations of guppies in nature. I created 80 replicate rearing groups per treatment, 40 with a focal male, and 40 with a focal female. To get individuals with the appropriate levels of *G* for these treatments, I used a six-generation-deep breeding design. Further information on this breeding design and the tank setup used for rearing focal individuals for this experiment can be found in the Supplementary Methods. Focal individuals were held in the rearing tanks alongside their group members until immediately before they were tested, as adults.

### Behavioral tests

I used behavioral assays to determine the patterns of kin discrimination exhibited by focal individuals. Because male guppies avoid pre-copulatory competition with closely related rivals, I conducted male-male competition trials to measure male kin discrimination. In my study population, male-male competition primarily consists of interruption behaviors (4). Interruptions occur when two males are pursuing a female simultaneously and the trailing male darts around the leading male, positioning himself directly behind the female. Guppies have internal fertilization and a male must position himself directly behind a female to attempt copulation (27, 42). Males that are interrupted more frequently have reduced mating opportunities (4), and male guppies use kin recognition to interrupt closely related rivals less frequently than non-kin rivals (4, 17). I therefore used differences in the number of times the focal male interrupted each rival as a measure of kin discrimination.

Female guppies behaviorally avoid inbreeding (30), so I conducted mate choice trials and used differences in female mating interest towards different males as a measure of kin discrimination. Females signal their mating interest in males by responding positively to male courtship displays (27) (see Supplementary Methods for descriptions of these behaviors). I measured female mating interest as the proportion of a male’s courtship displays to which the female responded positively.

For tests of both female mate choice and male-male competition, I observed groups of freely interacting fish in a large observation tank (90 x 45 x 37 cm), accommodating a wide range of guppy reproductive behaviors (27, 42). In each trial, I presented either a focal male or focal female with 5 non-focal males simultaneously: one each of *G* = 0.5, 0.375, 0.25, 0.125, and 0 to the focal individual. I chose, by eye, non-focal males that were similar to one another in their amounts of orange and black coloration, since these ornaments affect attractiveness in some guppy populations (27). I also included non-focal females in each trial (6 for mate choice trials, and 8 for intrasexual competition trials), to create female-biased sex ratios (males:females; 6:8 for intrasexual competition trials; 5:6 for mate choice trials) similar to those observed in natural guppy populations (28, 43). Non-focal females were unrelated (*G* = 0 within the breeding design) to all non-focal males and focal individuals used in the same trial. Furthermore, all non-focal individuals were unfamiliar with the focal individual because they were held in different tanks that were visually isolated from one another. This prevented effects such as individual recognition or a shared rearing environment from biasing the results. To eliminate any potential effects of familiarity acquired *in utero* or postnatally, any non-focal males that shared a mother with the focal individual (i.e. full-siblings, maternal sibling-cousins, and maternal half-siblings) were taken from the mother’s first brood, while the focal individual was taken from the second brood. Importantly, the non-focal individuals used in all treatments were reared under similar conditions, and spanned the same range of *G* values to the focal individual (as described above), irrespective of the focal individual’s treatment. Therefore, any between-treatment differences in the behavioral interactions I assayed can only be explained by the focal individual’s rearing environment.

Previous studies have shown that female guppies are relatively indiscriminate as virgins, but become choosy once mated (30). Therefore, I mated focal and non-focal females before using them in the behavioral trials, by pairing each female with a half-brother for 24 h prior to testing. The half-brothers used for these matings were not part of the focal female’s rearing group or testing group. I used half-brothers because *G* = 0.125 was close to the 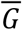 for all three treatments, which should have minimized any effect of exposure to these males on female population estimates. Because I paired females from all three treatments with half-brothers, the treatments were not confounded by differences in mating history.

To discourage foraging behavior during behavioral trials, I fed focal and non-focal individuals 1 h before testing. I placed the focal male in the observation tank and, after a 15-min acclimation period, added the non-focal fish and started the trial. I recorded the ensuing interactions for 30 min using JWatcher™ (version 1.0, available at http://www.jwatcher.ucla.edu). To account for diel variation, all trials were performed between 9:00 and 11:30 AM, when guppies are sexually active (27). Between each trial, I performed a partial water change and ran a carbon filter in the observation tank for at least 30 min to minimize any carry-over of olfactory cues. Trials were performed in a random order with respect to treatment, meaning that any such carry-over should not have confounded the results. I carried out all observations and was blind to the focal individual’s treatment and the consanguinity of the non-focal males at the time of testing.

### Statistical analyses

To determine whether kin discrimination is affected by the phenotypic distribution of the social environment in which an individual develops, I asked whether the behaviors of interest (mating interest or numbers of interruptions) were significantly affected by non-focal male *G*, rearing treatment, or their interaction. I used permutational ANOVA (*R* v3.3.3, *Adonis* function in package *vegan* v2.4-2), which calculates the pairwise distances among data points, and then performs a permutation test (randomly assigning labels amongst factors) to partition the distance matrix among sources of variation. Pseudo-F ratios are then used to compute *P* values. Each focal individual’s behavior towards a given non-focal male constitutes a different observation in the dataset, resulting in the high degrees of freedom (see results). Unlike parametric ANOVA, permutational ANOVA does not assume normality, homogeneity of variances, or independence among observations. In cases of non-independence, pseudo-replication can be avoided by accounting for the covariance structure of the data. In this experiment, non-independence occurred because *G* varied within trials (i.e. among the 5 non-focal males with whom the focal individual interacted). I accounted for this within-subjects design in the model by defining the identity of the focal individual as strata nested within treatment. Analogous to creating a blocking factor (44), this constrained permutations among levels of *G* to within each trial, thus building the within-subjects covariance structure into the null distribution used for hypothesis testing. I performed two analyses for each sex: one comparing the control treatment against the high similarity treatment, and another comparing the control treatment against the high variance treatment.

I performed post-hoc tests as follows. In cases where the interaction between treatment and non-focal male *G* was significant, I performed two-sample permutational t-tests (package *DAAG* v1.24) examining the simple effect of treatment within each level of non-focal male *G*. In cases where I observed a significant main effect of *G*, I performed contrasts examining the change in behavior with each marginal change in the level of *G*. This allowed me to ask, for each treatment, whether focal individuals discriminated based on fine-scale differences in *G.* I accounted for the non-independence of observations taken on the same focal individual in the same trial by using a single, composite measure for each focal individual’s change in behavior across the marginal change in *G*. This composite measure was the difference in behavior between the two levels of *G* being compared (e.g. number of interruptions towards *G* = 0 minus number of interruptions towards *G* = 0.125). I then used one-sample permutational t-tests (package *DAAG* v1.24) to ask whether the distribution of these difference values was significantly different from 0, which is the null expectation in the absence of kin discrimination. I performed the planned contrasts separately for each treatment (even in cases where the interaction between *G* and treatment was not significant) so that the sample sizes would be equal. This facilitated meaningful comparison of the results of these planned contrasts among treatments. To prevent multiple comparisons from increasing the type-1 error rate, I applied the Bonferroni-Holm correction to each set of planned and post-hoc tests. This method offers the same level of protection against type-1 error as the Bonferroni method but retains greater statistical power, and is valid under broader assumptions than the Hochberg approach (45). I report both the raw and Bonferroni-Holm corrected *P* values. For all permutational ANOVA, I used Euclidean distance matrices. For each permutational test (t-tests and ANOVA) I calculated *P* values using 10 000 simulations. All analyses were performed in R v3.6.3.

## Results

### Male-male competition

Comparing the high similarity and control treatments, I found that the number of interruptions focal males directed towards their competitors was significantly affected by *G* (pseudo-*F*_1,396_ = 254.310, *P* = 0.001) and rearing treatment (pseudo-*F*_1,396_ = 19.193, *P* = 0.001), but not their interaction (pseudo-*F*_1,396_ = 0.112, *P* = 0.751). Thus, males directed more competitive effort towards rivals of greater *G*, and performed more competitive behaviors overall if they were reared in the high similarity treatment (Figure 3A). Consistent with my predictions, this suggests that focal males in the high similarity treatment perceived lower levels of relatedness to their rivals than did focal males from the control treatment. Planned contrasts among levels of *G* revealed, in both the high similarity and high variance treatments, significant reductions in the numbers of interruptions with every marginal increase in the level of *G* (Table 1A). Focal males in the control and high similarity treatments therefore discriminated at a fine-scale among rivals of different levels of *G*.

**Table 1.** Results of one-sample permutational t-tests within treatments, testing the behavior of focal individuals was affected by marginal changes in consanguinity (*G*).

| | Comparison ($G$) | $P^*$ | $P^*$ (corrected<sup>†</sup>) |
| --- | --- | --- | --- |
| a. Interruptions |  |  |  |
| Control | 0 vs. 0.125 | **0.024** | **0.048** |
|  | 0.125 vs. 0.25 | **0.002** | **0.008** |
|  | 0.25 vs. 0.375 | **0.012** | **0.036** |
|  | 0.375 vs. 0.5 | **0.049** | **0.049** |
| High similarity | 0 vs. 0.125 | **0.048** | **0.048** |
|  | 0.125 vs. 0.25 | **0.022** | **0.048** |
|  | 0.25 vs. 0.375 | **0.015** | **0.048** |
|  | 0.375 vs. 0.5 | **0.012** | **0.048** |
| High variance | 0 vs. 0.125 | 0.984 | 0.984 |
|  | 0.125 vs. 0.25 | **0.045** | 0.135 |
|  | 0.25 vs. 0.375 | **0.008** | **0.032** |
|  | 0.375 vs. 0.5 | 0.416 | 0.832 |
| b. Female mating interest |  |  |  |
| Control | 0 vs. 0.125 | 0.115 | 0.115 |
|  | 0.125 vs. 0.25 | **0.023** | **0.046** |
|  | 0.25 vs. 0.375 | **0.015** | **0.045** |
|  | 0.375 vs. 0.5 | **0.001** | **0.004** |
| High similarity | 0 vs. 0.125 | **0.016** | **0.048** |
|  | 0.125 vs. 0.25 | **0.021** | **0.048** |
|  | 0.25 vs. 0.375 | **0.022** | **0.048** |
|  | 0.375 vs. 0.5 | **0.003** | **0.012** |
| High variance | 0 vs. 0.125 | 0.504 | 1.000 |
|  | 0.125 vs. 0.25 | **0.027** | 0.100 |
|  | 0.25 vs. 0.375 | **0.025** | 0.100 |
|  | 0.375 vs. 0.5 | 0.740 | 1.000 |

\*Significant $P$ values are bolded.
<sup>†</sup> $P$ values after applying the Bonferroni-Holm correction.

**Figure 3.**
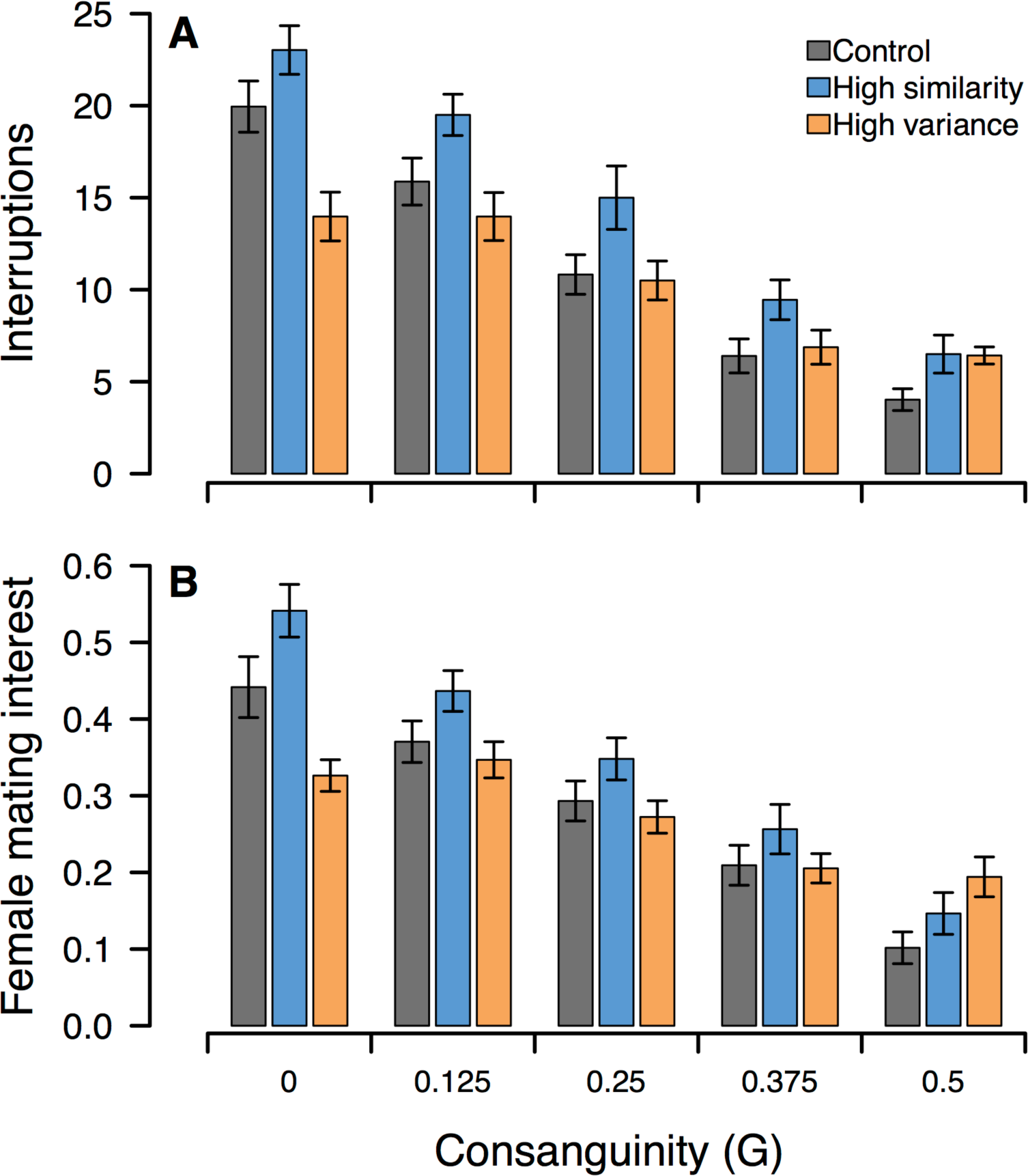
Numbers of interruptions performed by focal males (A), and mating interest of focal females (B), towards males of different consanguinities. Error bars represent mean ± s.e.m; n = 40 focal individuals per treatment per panel; each focal individual was tested with one non-focal male of each level of G.

Comparing the high variance and control treatments, I detected a significant main effect of *G* (pseudo-*F*_1,396_ = 174.384, *P* = 0.001) and a significant interaction between *G* and treatment (pseudo-*F*_1,396_ = 15.806, *P* = 0.002), while there was no significant main effect of treatment (pseudo-*F*_1,396_ = 2.451, *P* = 0.131). Therefore, as expected, males from the high variance treatment exhibited weaker (i.e. less extreme) kin discrimination towards rival males of different *G* (Figure 3A). Planned contrasts among levels of *G* revealed that focal males from the high variance treatment directed significantly more interruptions towards males of *G* = 0.25 than *G* = 0.375 (*P* = 0.032), but I did not find significant discrimination across other marginal changes in the level of *G* (Table 1A). These results suggest that focal males from the high variance treatment did not exhibit the same degree of fine-scale kin discrimination observed in the control and high variance treatments, providing further support for the prediction males reared in a social group with greater phenotypic variation should be less discriminatory. Comparing numbers of interruptions between the control and high variance treatments within each level of *G*, I found significant differences at *G* = 0 and *G* = 0.5, but not at other levels of *G* (Table 2A). Therefore, focal males from the high variance and control treatments behaved similarly towards rivals of intermediate *G* values, but their behaviors towards rivals with extreme (high or low) *G* values differed. This result further supports the prediction that individuals from the high variance treatment have the same average perception of relatedness to conspecifics as control individuals, but exhibit less extreme discrimination based on differences in rival *G*.

**Table 2.** Results of two-sample permutational t-tests within each level of *G*, testing whether the behavior of focal individuals differed among treatments.

| | $G$ | $P^*$ | $P^*$ (corrected<sup>†</sup>) |
| --- | --- | --- | --- |
| a. Interruptions |  |  |  |
| Control vs. high variance | 0 | **0.007** | **0.028** |
|  | 0.125 | 0.263 | 0.789 |
|  | 0.25 | 0.741 | 1.000 |
|  | 0.375 | 0.861 | 1.000 |
|  | 0.5 | **0.003** | **0.015** |
| b. Female mating interest |  |  |  |
| Control vs. high variance | 0 | **0.012** | **0.048** |
|  | 0.125 | 0.514 | 1.000 |
|  | 0.25 | 0.532 | 1.000 |
|  | 0.375 | 0.904 | 1.000 |
|  | 0.5 | **0.007** | **0.035** |

\*Significant $P$ values are bolded.
<sup>†</sup> $P$ values after applying the Bonferroni-Holm correction.

### Female preference

Comparing the high similarity and control treatments, I observed an effect of *G* (pseudo-*F*_1,396_ = 194.9, *P* = 0.001) and treatment (pseudo-*F*_1,396_ *=* 11.609, *P* = 0.001) on female interest, while their interaction was not significant (pseudo-*F*_1,396_ = 0.985, *P* = 0.325). Thus, consistent with predictions, females displayed lower mating interest towards males of higher *G*, and exhibited higher mating interest overall if reared in the high similarity treatment (Figure 3B). Females from the control treatment exhibited significantly less mating interest with each marginal increase in the level of *G*, except that females did not discriminate significantly between males of *G* = 0 and *G* = 0.125 (Table 1B). In the high similarity treatment, female interest declined significantly with each marginal increase in *G* (Table 1B). Focal females in the control and high similarity treatments therefore discriminated at a fine-scale among rivals of most different levels of *G*.

Comparing the high variance and control treatments, I found an effect of *G* (pseudo-*F*_1,396_ = 118.674, *P* = 0.001), no significant main effect of treatment (pseudo-*F*_1,396_ = 0.776 *P* = 0.384), and a significant interaction between *G* and treatment (pseudo-*F*_1,396_ = 14.464, *P* = 0.001). Thus, females from the high variance treatment were less discriminatory among males of differing levels of *G* (Figure 3B). Planned contrasts did not detect any significant differences in female mating interest across any marginal changes in *G* (table 1B), further indicating that females from the high variance treatment exhibited less extreme kin discrimination compared to the control and high similarity treatments. Post-hoc tests comparing female mating interest between the control and high variance treatments within each level of *G* revealed significant differences at *G* = 0 and *G* = 0.5, but not at other levels of *G* (Table 2B). Focal females from the control and high variance treatments therefore behaved similarly towards rivals of intermediate *G* values, but differed in their behavior towards rivals with extreme *G* values.

## Discussion

To assess relatedness to a given conspecific based on phenotypic similarity, it is widely assumed that an evaluator uses information about (i) its own phenotype, and (ii) the phenotype of the given conspecific (13, 14). However, recent theory indicates that this model of phenotype matching is incomplete; for kin recognition to be reliable, evaluators must also consider information about the population distribution of phenotypes (21). I have demonstrated that varying the genotypic (and hence, phenotypic) composition of an individual’s rearing group affects its patterns of kin discrimination in the manner predicted by population estimation theory. These results provide the first empirical evidence suggesting that population estimates inform phenotype matching, and have dramatic effects on mate choice and intrasexual competition. Focal individuals in all three treatments were tested with equivalent sets of conspecifics, and developed in similar environments, except for the differences in the social groups within which they developed. The behavioral differences that I observed among treatments can therefore only be explained by individuals encoding information related to the phenotypic distribution of their social group and adjusting their patterns of behavioral discrimination accordingly. Thus, the results show that plasticity contributes to population estimates, and that this plasticity is mediated by social experience. Collectively, these findings indicate that individuals recognize kin by assessing their phenotypic similarity to conspecifics, relative to their social group.

I found that focal individuals adjusted their behavior in a graded manner in response to differences in *G* to non-focal males, indicating that guppies are capable of fine-scale kin discrimination. As expected, individuals from the high similarity treatment, who were more genetically (and hence, phenotypically) similar to members of their social group than those in the control treatment, treated novel conspecifics as though they were less closely related. This is consistent with the prediction that individuals form a population estimate based on the phenotypes of their group members, such that individuals that are more similar to their group members perceive conspecifics to less similar, and hence, less closely related, relative to their population estimate. Additionally, I found that rearing individuals within social groups that were more genetically (and hence, phenotypically) variable caused them to exhibit less extreme discrimination among conspecifics of different levels of consanguinity. This result is consistent with the prediction that individuals incorporate information about phenotypic variance into their population estimates, such that individuals from more variable groups acquire “wider” population estimates, making differences in phenotype appear less extreme. Thus, guppy kin recognition is informed by population estimates that reflect both the mean and variance of the phenotypes in their rearing groups. Furthermore, population estimation appears to play a similar role in kin recognition for both sexes, and in the contexts of both inbreeding avoidance and nepotism during intrasexual competition.

It is not clear what specific physiological and/or neurological processes underpin the formation or use of population estimates. Guppies have been shown to recognize individuals within groups of several dozen individuals (46, 47), indicating they are capable of storing phenotypic information about large numbers of conspecifics to form population estimates. It is also possible that guppies may use heuristics that reduce the cognitive demands of forming population estimates. One possible heuristic could consist of individuals encoding a “similarity scale” centered on the population mean phenotype, bounded on one side by the kin template, and on the other side by the most dissimilar phenotype observed (21). The evaluator should perceive a conspecific to be positively related to the degree that its phenotype is closer to the kin template than the population average, or negatively related to the extent that its phenotype is more distant than the population average. This heuristic offers a simple, yet general process by which organisms could distinguish between positive and negative relatives. Future work should investigate this heuristic model, and mechanism(s) by which population estimates are stored (e.g. long-term memory and/or changes in the sensory peripheries).

Collectively, these findings provide fundamental insight into the proximate mechanisms underpinning phenotype matching: when comparing their own phenotype against that of a conspecific, individuals use population estimates to contextualize this comparison within the “phenotypic space” of their social group. That these population estimates are plastic should allow guppies to respond rapidly to variation in the population distribution of phenotypes over time and/or space. As variation in phenotypic distributions across time and space is prevalent in nature in many taxa, I suggest that plastic population estimation may be a common property of phenotype matching systems.

In the kin selection literature, numerous theoretical and empirical studies have emphasized the importance of scale in determining the inclusive fitness consequences of social actions. Specifically, kin selection favours actions that increase the fitness of conspecifics that are positively related (i.e. *G* greater than 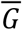) at the scale at which social interactions and their fitness consequences play out–sometimes called the scale of competition, or the interaction neighborhood (34, 48–50). Consequently, the scale of competition is a key factor shaping the evolution of nepotism, spite, and cooperation (38). Population estimation provides a critical link between kin recognition and the scale of competition by calibrating phenotype matching to the “phenotypic space” of the evaluator’s social group, which for many kinds of social interactions should approximate the scale of competition. For this reason, population estimation should enhance phenotype matching, thereby increasing opportunities for the evolution of nepotism, inbreeding avoidance, and other social strategies that are modulated by kin recognition.

Krupp and Taylor (21) note patterns of kin recognition in several other species match the predictions of population estimation theory. For example, argentine ants from colonies with greater phenotype variability are less aggressive towards foreign conspecifics (9). This is believed to modulate the extent to which different colonies co-exist or competitively exclude one another, driving large-scale differences in invasiveness and population structure (51). Additional studies have documented patterns of behavioral discrimination that are suggestive of population estimation in social ants (7, 52, 53), the Columbian ground squirrel (54), the great reed warbler (55), and humans (56). Alternative explanations for each of these examples may exist (57, 58); nevertheless, in combination with my results these examples suggest that population estimation may be taxonomically widespread, with consequences for a variety of ecologically and evolutionarily important social interactions.

That plastic population estimates inform phenotype matching has important implications for the design and interpretation of kin recognition studies, particularly in terms of ecological relevance. Some hundreds of laboratory experiments have tested for phenotype matching and explored details of the underlying mechanisms, in the guppy and many other taxa (13, 14). These studies have made valuable contributions to our understanding of kin recognition, but focal individuals in these studies are often reared within social groups that differ drastically in their distribution of *G* from what the organisms would encounter in nature. For example, lab stocks are often inbred relative to source populations and are held in tightly controlled environments, meaning that the total genetic and environmental variation in phenotype tends to be lower. Following our results, such conditions would be expected to lead individuals to form population estimates that are “narrow” and highly similar to themselves, making even subtle phenotypic differences have an unnaturally large impact on relatedness judgments. In this way, population estimation could help to explain why lab studies have implicated particular phenotypic cues, such as olfactory components associated with the major histocompatibility complex (MHC) (59–62) or the major urinary protein gene cluster (MUP) (63, 64), as being important for kin recognition, while evidence that these same cues are important in natural settings remains somewhat limited (59, 61, 62). If population estimation is taxonomically widespread, tests that incorporate natural genetic and environmental variation (62, 65), and ecologically-relevant social settings, will be especially important for clarifying the generality of lab-based findings to natural contexts.

I suggest that population estimates may be used by organisms in behavioral contexts beyond kin recognition. Altering the distribution of color pattern phenotypes of the social group within which female guppies develop can affect their mating preferences (66–68). The effects observed in these studies were complex and differed from the patterns I report here. Nevertheless, they demonstrate that female guppies encode information about the phenotypic distributions of (male) conspecifics and alter their mating decisions in ways that do not appear related to kin recognition. There is also reason to think that population estimates may be important in species recognition. Many animal taxa discriminate conspecifics from heterospecifics in social contexts including aggression, territoriality and mate choice. While some aspects of the underlying mechanisms remain poorly understood, there is evidence for both social and genetic influences on so-called species recognition templates (69–72). A population estimate representing the mean and variance in phenotype could be useful as a species template, as a conspecific is more likely to fall within the population’s phenotypic distribution than a heterospecific. Determining the extent of potential overlap in the mechanisms used to sample the population distribution of phenotypes for kin recognition and for other social interactions could help shed light on the evolutionary pressures shaping population estimation.

In summary, I have provided experimental evidence that plastic population estimates inform phenotype matching in the guppy, with dramatic consequences for social behaviors. These results support the first complete model of phenotype matching, providing fundamental insight into the proximate mechanisms that make this type of kin recognition reliable: individuals are able to assess relatedness to conspecifics by assessing how phenotypically similar a given conspecific is relative to the social group. My results demonstrate two important consequences of population estimation for social behavior. All else being equal, individuals that are more similar to their social group tend to treat conspecifics as less closely related, and individuals from more diverse social groups will tend to exhibit weaker kin discrimination. I found that population estimation was used by both sexes, in the contexts of nepotism and inbreeding avoidance. By allowing individuals to reliably infer relatedness from phenotypic information, population estimation increases opportunities for the evolution of these and other behavioral strategies that are conditioned on kinship. Data from additional taxa and behavioral contexts hint that population estimation may be widespread, and thus of broad importance for understanding social evolution. Future work should therefore investigate the generality of this mechanism.

## Acknowledgements

I am grateful to F.H. Rodd, L. Rowe, D. McLennan, A. Wardlaw, R. Williamson, D. Queller, A. Agrawal, A. De Serrano, A. Li, M. Foisy, J. Levine, K.A. Hughes, A.M. Makowicz, and J.J. Valvo for insightful comments and discussion. I also thank I. Ramnarine and the Government of Trinidad and Tobago for permission to collect the guppies used to establish the lab population. Many undergraduates helped to maintain the lab population. This work was supported by funding from the National Sciences and Engineering Research Council (NSERC) to M.J.D (grant no. CGSM-427283-2012).

