## Supplementary Materials for "It’s all relative: population estimates enhance kin recognition in the guppy"

Mitchel J. Daniel

**Supplementary Materials**

### Supplementary Methods

*Breeding crosses*

I generated the individuals used in the rearing groups by performing sets of six-generation-deep breeding crosses. The first generation consisted of adults drawn from stock tanks, each mated to an individual from a different tank to avoid inbreeding. There was no inbreeding within the breeding design, meaning that the results cannot be explained by differences in inbreeding coefficients. I drew focal individuals and their group members from the fifth and sixth generations. To avoid pseudoreplication, all focal individuals were no more consanguineous to one another than second cousins ($G$ = 0.0313). Here, I describe the relationships between the focal individuals from each treatment and the non-focal individuals used as group members or in behavioral tests (depicted in Supplementary Figure S1).

In the control treatment, the focal individual’s group members consisted of four uncle/aunt-cousins once removed (i.e. offspring produced by crossing one of the focal individual’s grand-parents with a full-sibling of the focal individual’s other grand-parent; $G$ = 0.188), eight half-uncles/aunts (i.e. half-siblings of one of the focal individual’s parents; $G$ = 0.125), and four first cousins once removed ($G$ = 0.063). In the control treatment, all group members that had the same type of relationship (e.g. all four first cousins) to the focal individual were produced by the same breeding pair. Focal individuals from the control treatment were tested with one full-sibling ($G$ = 0.5), one sibling-cousin (i.e. offspring produced by crossing one of the focal individual’s parents with a full-sibling of the focal individual’s other parent; $G$ =

0.375), one uncle ($G$ = 0.25), one half-uncle, and one non-kin male (i.e. a male that is unrelated to the focal individual within the breeding design; $G$ < 0.002).

In the high-similarity treatment, the focal individual's group members consisted of four sibling-cousins, eight half-siblings ($G$ = 0.25), and four first cousins ($G$ = 0.125). As in the control group, all group members that had the same type of relationship to the focal individual were produced by the same breeding pair. Group members for the high-similarity treatment were derived from the same crosses within the breeding design as for the control treatment. Consequently, group members had the same relationships to one another as in the control treatment. The critical difference between the control and high similarity treatments it that the focal individual was produced one generation earlier in the breeding design for the high similarity treatment, which doubled the focal individual's consanguinity to group members. Focal individuals from the high-similarity treatment were tested with one full-sibling, one sibling-cousin, one half-sibling, one first cousin, and one non-kin male.

In the high-variance treatment, the focal individual's group members consisted of four nephew/niece-cousins once removed (i.e. offspring produced by a full-sibling of the focal individual crossed with a first cousin of the focal individual; $G$ = 0.1875), eight half-nephew/nieces (i.e. offspring of the focal individual's half-sibling; $G$ = 0.125), and four first cousins once removed. In the high-variance treatment, group members that had the same type of relationship to the focal individual were cousins of one another. Furthermore, they were derived from one generation later in the breeding design than the group members used in the control treatment, while the focal individual was derived from one generation earlier in the breeding design than the control treatment (Supplementary Figure S1). These differences in breeding design quartered consanguinity among group members without altering the focal individual's consanguinity to the group members. Focal individuals from the high-variance treatment were tested with one full-sibling, one sibling-cousin, one nephew ($G$ = 0.25), one half-nephew, and one non-kin male.

For some of the relationships in this breeding design, we had to decide whether to make individuals related through a female or a male for one or more of the crosses involved (e.g. maternal first cousins vs. paternal first cousins). For each such cross, we randomized whether individuals were related through the maternal or paternal line. Consequently, parent-of-origin effect and maternal or paternal effects should not have biased phenotypic similarity, or social behaviors, among treatments or levels of non-focal male $G$. Exception to this randomization was made for cases in which a single individual had to be crossed with multiple mates to produce the necessary relationships (e.g. half-siblings). We re-mated males rather than females that parentage was unambiguous.

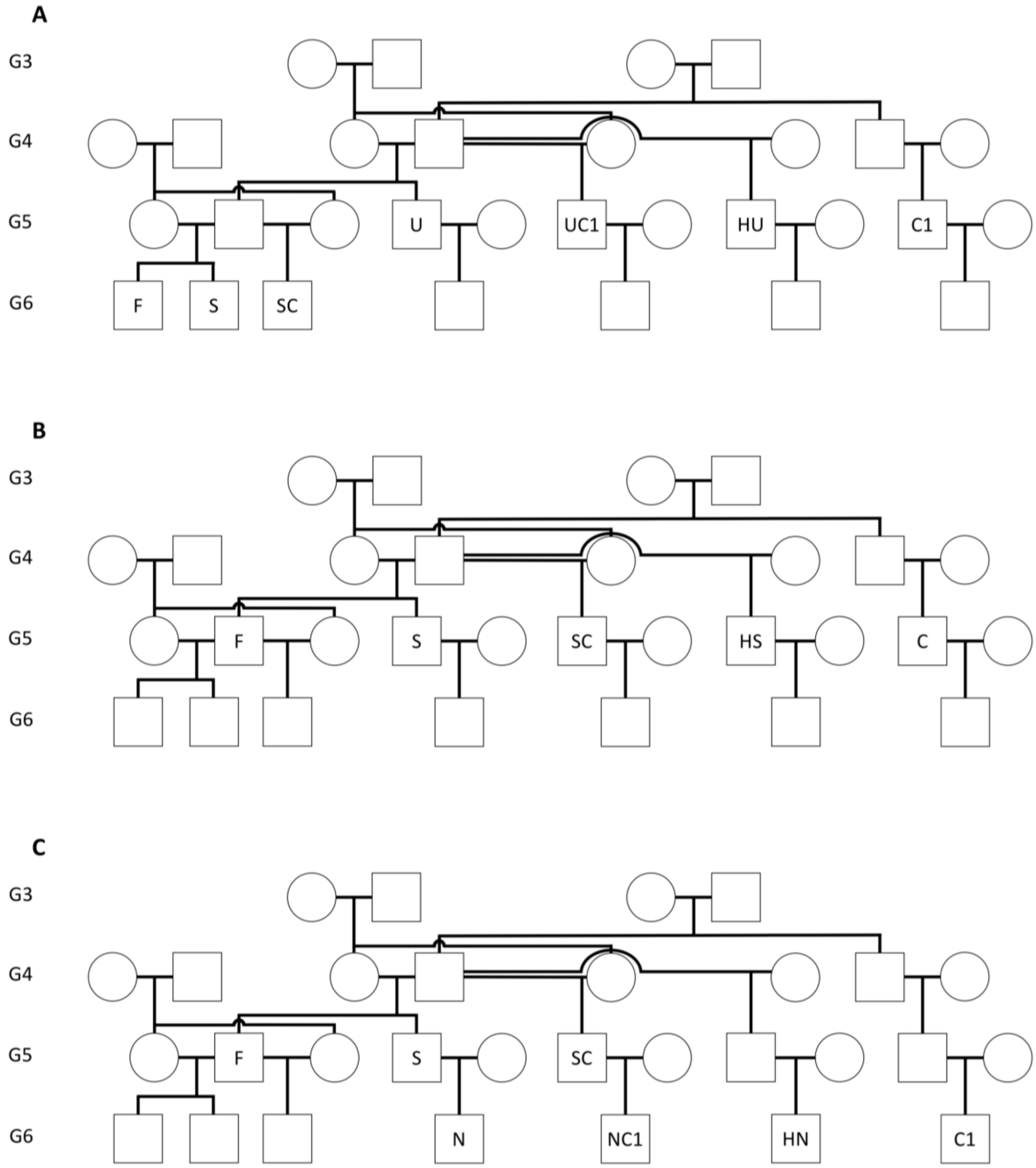


Supplementary Figure S1. Pedigree depicting the types of relationships between the focal individual, the members of its rearing group, and the non-focal males used in behavioral tests for (*A*) the control treatment, (*B*) the high similarity treatment, and (*C*) the high variance treatment. This pedigree spans generations three through six (G3-6) of the breeding design. Individuals do not have kinship within the breeding design except through the relationships depicted in this pedigree. Relationships are denoted as follows: F = focal individual, S = full sibling, SC = sibling-cousin, HU = half-uncle/aunt, U = uncle/aunt, UC1 = uncle/aunt-cousin once removed, C1 = cousin once removed, HS = half-sibling, C = cousin, HN = half-nephew/niece, N = nephew/niece, NC1 = nephew/niece-cousin once removed. Non-kin are not displayed on the pedigree. The sexes (circles denote females; squares denote males) shown on this pedigree are arbitrary.

*Husbandry and rearing setup*

Focal individuals were placed into 20 L rearing tanks (40 x 25 x 20 cm) with their assigned group members within 24 h after birth. Focal individuals were born up to one month after all of their group members, so that they were of similar age to their group members, and so that all group members were present when the focal individual was added to the rearing tank. In nature, juvenile guppies are highly philopatric and, shortly after birth, begin shoaling with conspecifics of similar age (1, 2). Therefore, this setup simulates natural association among similarly aged conspecifics.

Each rearing tank included an internal divider that separated the focal individual from its group members. This prevented the focal individual from mating (which might have biased subsequent behavior), and allowed me to distinguish the focal individual from the other fish. These dividers were Plexiglas with a mesh insert, permitting both visual and olfactory communication. Guppies are able to establish individual recognition and recognize kin across these dividers (3, 4), so this setup should not have impeded focal individuals from assessing group member phenotypes. The tanks were lined, on the bottom, with neutral colored gravel and lit by full-spectrum fluorescent bulbs. I fed the fish Tetramin© fish food and live nauplii larvae of *Artemia salina* twice daily, and maintained a 12:12 h light:dark cycle, at 25-26° C. Guppies mature at approximately 100 days old. I held focal individuals and their group members in this setup until immediately before testing, at 120 to 140 days old.

*Behavioral protocols*

Guppies exhibit a promiscuous mating system with internal fertilization. Male guppies are persistent in their mating efforts and flexibly exhibit two mating strategies. The first of these is sneak mating attempts, in which the male thrusts his gonopodium at the female's gonopore without solicitation (1, 5). The second male mating strategy is courtship, in which the male positions himself within the female's field of view and quivers with an arched back (1). The female may exhibit mating interest by orienting towards the displaying male, approaching, and occasionally performing a "glide" response (1, 6). We measured the mating interest of the focal female towards a given non-focal male as the proportion of his courtship displays to which she responded positively with one or more of these behaviors. Copulations rarely occur within the timeframe of mating trials, and were thus too sparse to analyze. However, female positive responses are strongly predictive of male mating success (1, 7), and are thus a reliable indicator of female mate choice. Orienting, approaching, and gliding behaviors have been widely used to measure female mating preferences in this species (7–14), including in studies of inbreeding avoidance (3).

To access the female's gonopore for a coercive or cooperative mating attempt, a male must position himself directly behind the female. When two males simultaneously pursue the same female, they frequently jockey for position behind the female. This involves interruption behavior, in which the trailing male darts around the leading male, usurping access to the female (15, 16). We recorded the number of times the focal male interrupted each of the non-focal males. Because males direct fewer interruptions towards more closely related rivals (15), we used the number of times the focal male interrupted a given rival to infer perceived relatedness.
